# Can mechanistic constraints on recombination reestablishment explain the long-term maintenance of degenerate sex chromosomes?

**DOI:** 10.1101/2023.02.17.528909

**Authors:** Thomas Lenormand, Denis Roze

## Abstract

Y and W chromosomes often stop recombining and degenerate. Most work on recombination suppression has focused on the mechanisms favoring recombination arrest in the short term. Yet, the long-term maintenance of recombination suppression is critical to evolving degenerate sex chromosomes. This long-term maintenance has been little investigated. In the long term, recombination suppression may be maintained for selective reasons (e.g., involving the emergence of nascent dosage compensation), or due to mechanistic constraints preventing the reestablishment of recombination, for instance when complex chromosomal rearrangements evolve on the Y. In this paper, we investigate these ‘constraint’ theories. We show that they face a series of theoretical difficulties: they are not robust to extremely low rates of recombination restoration; they would rather cause population extinction than Y degeneration; they are less efficient at producing a non-recombining and degenerate Y than scenarios adding a selective pressure against recombination, whatever the rate of recombination restoration. Finally, whether such very high constraints exist is questionable. Very low rates of recombination reestablishment are sufficient to prevent Y degeneration, given the large fitness advantage to recover a non-degenerate Y or W for the heterogametic sex. The assumption of a lack of genetic variation to restore recombination seems also implausible given known mechanisms to restore a recombining pair of sex chromosomes.

## Introduction

How could the degeneration of whole Y or W chromosomes evolve without being opposed by natural selection? Several theories have been put forward to explain this problem, often separating it into several subproblems: Why does recombination initially stop between sex chromosomes? Why does degeneration occur on a nonrecombining part of a genome? Why is recombination suppression maintained in the long term, once degeneration has already caused substantial degradation of the Y or W? Why isn’t recombination eventually reestablished to limit maladaptation? When considering the suppression of recombination, it is useful to distinguish between the initial arrest and its long-term maintenance. We first briefly mention the different ideas that have been proposed for the initial arrest, which has attracted the most attention. We then present the different mechanisms that could lead to the long-term maintenance of recombination suppression, which has been much less investigated. We discuss the XY case throughout the paper, but all arguments apply to ZW systems as well.

### The initial recombination arrest on sex chromosomes

Six main ideas have been proposed to explain short-term recombination arrest on sex chromosomes. Even if we focus on the long-term maintenance of recombination suppression, it is useful to list these ideas to clarify the difference between the short and long-term suppression of recombination.

The first proposes that the sex determination (SD) locus happened to arise in a region of the genome where recombination was already suppressed [1,2] or in a non-recombining genome. In achiasmate species, in particular, the heterogametic sex does not recombine (the so-called Haldane-Huxley rule [3,4]), so that, wherever the SD locus is located, the Y or W will not recombine. The direction of causality between the evolution of suppressed recombination on sex chromosomes and the evolution of achiasmy is however difficult to establish [4].

The second idea proposes that recombination is selected against to prevent the production of neuter individuals in species where sex is determined by a combination of a male-sterility and a female-sterility locus [5,6]. This explanation applies to species with such specific combinations of SD loci and is likely to apply to the evolution of separate sexes from dioecy or genetic sex determination from environmental sex determination. However, it cannot account for later events of recombination suppression beyond the region of the genome containing the SD loci.

The third idea is the “sexually antagonistic selection” scenario [6–12], where suppression of recombination is selectively favored due to the occurrence of loci with sexually antagonistic (SA) effects on sex chromosomes. In this case, a non-recombining Y benefits from a selective advantage, by permanently combining male-determining and male-beneficial alleles. A variant of this scenario involves sex differences in the intensity of selection (but not in the direction of selection as considered in SA theory) against deleterious mutations [13]. There is plenty of evidence for SA variation [14,15], making this hypothesis very plausible. However little evidence for this scenario has been obtained despite intensive research [1,10,16], partly due to the difficulty of definitively establishing the role of SA variation in driving recombination suppression.

The fourth scenario involves ”lucky” inversions on the Y capturing the SD locus [17–19]. Inversions capture at birth a sample of segregating deleterious variation. Some “lucky” inversions can have a selective advantage because they initially capture a portion of the proto-Y that carries fewer or milder deleterious mutations, compared to the population average. Their initial fitness advantage decays through time but can be sufficient to allow them to fix [17,18]. Much of this decay is caused by a deterministic return to the population average load [18,20], but selective interference also contributes. This decay rate is larger for larger inversions [17,18], but inversions with a large initial advantage are likely to be sampled among larger inversions [17]. These opposing effects determine the distribution of fixed inversion sizes, with a mode biased toward smaller sizes compared to their input size distribution [17,18]. Contrary to the fixation of ‘lucky’ inversions on autosomes [20], the fixation of inversions capturing the SD locus on the proto-Y is expected to cause recombination suppression on the inverted segment of the Y [16,21]. This process can involve any Y variant suppressing recombination around the SD locus and closely linked to this region, not only inversions (we use the term inversion only for simplifying the presentation). For instance, it could involve changes in chromatin structure or the loss of recombination hotspots. It solely depends on the presence of deleterious mutations and variation in recombination rates and is thus expected to be widespread. However, due to its recent description, it has not been investigated empirically.

The fifth mechanism involves recombination suppression near the sex-determining locus due to the neutral accumulation of sequence divergence that decreases homology between the X and Y and suppresses recombination as a side effect [22]. When a small non-recombining region is established, then this accumulation continues by slowly moving the boundary of the pseudo-autosomal region. The idea is that strict homology is required for recombination to occur. However, while this effect has been documented in mitotic lineages [23–25, and other references cited in 22], it is not clear whether small amounts of divergence prevent recombination during meiosis. In particular, data from tetraploid rye and from crosses among strains of *Arabidopsis thaliana* with varying levels of divergence suggest that heterozygosity may enhance rather than inhibit crossovers [26–28].

The last mechanism proposes that XY recombination suppression is favored because it increases heterozygosity around the SD locus in the heterogametic sex. Recent versions of this idea proposed that tight linkage between the SD locus and overdominant mutations, or combinations of recessive deleterious mutations (generating pseudo-overdominance) would be favored because these mutations would then be more often present in the heterozygous state. In partially inbred populations, linkage to the SD locus indeed increases heterozygosity, and this can favor recombination suppression in the presence of overdominant mutations [29], while more work is needed for the case of recessive deleterious mutations and random mating [18].

### The long-term maintenance of recombination arrest on sex chromosomes

It has been long established that in the absence of recombination, deleterious mutations will tend to accumulate on the Y due to selective interference [11,12,30–35], which should generate some maladaptation, especially in males. Why then is recombination not reestablished when degeneration becomes too strong? Three main ideas can be distinguished.

The first is that SA effects are sufficiently strong to maintain recombination arrest despite the accumulation of deleterious mutations. This has nevertheless been questioned [36–38]. Indeed, the deleterious effect of Y degeneration may eventually offset the selective advantage of linking male-determining and male-beneficial alleles. The restoration of recombination, if it is possible, may then become favorable. However, new SA mutations may continue to appear and accumulate on sex chromosomes, giving time for the population to evolve Y silencing / dosage compensation (DC) limiting maladaptation caused by Y degeneration.

The second idea is based on regulatory evolution. Once recombination is suppressed, cis regulators of gene expression may diverge between the X and Y, for genes located in the non-recombining portion of the Y. This regulatory instability may lead to the evolution of early DC (through the joint evolution of cis and trans-acting factors), concomitant with Y early silencing and degeneration. In the model proposed by [17], the emergence of this DC builds up pervasive SA regulatory effects, selectively preventing the long-term reestablishment of recombination. Note that regulatory evolution may lead to the accumulation of deleterious mutations and degeneration even in conditions where selective interference is inoperative [39].

The third possibility, which we term the “constraint” scenario, involves cases where recombination suppression is maintained in the long term despite the fact that it has become disadvantageous, due to mechanistic constraints preventing the reestablishment of recombination. Most models for the evolution of sex chromosomes ignore the possibility that recombination can be reestablished, implicitly assuming that a constraint maintains recombination arrest in the long term. Few models present a more detailed reasoning about this constraint. For instance, in Jeffries et al.’s simulation model of neutral arrest of recombination [22], crossovers are assumed to be fully suppressed once sequence divergence becomes too high. The constraint emerges from the loss of homology. However, the authors note that, in reality, rare recombination events could occasionally occur at high sequence divergence. While Jeffries et al.’s model does not include deleterious mutations, degeneration would generate strong selection to restore recombination and these rare events would be highly beneficial. Another idea is that reestablishing recombination might be difficult once complex chromosomal rearrangements have occurred on the Y. For instance, Jay et al.’s model [19] assumes that recombination is suppressed as soon as an inversion occurs and that the occurrence of secondary inversions (overlapping or occurring within a first one) prevents reversion of the first one. Hence, recombination could only be reestablished in a region if all inversions are exactly reinversed, irrespectively of the actual collinearity (or lack thereof) between the X and Y. This can allow lucky inversions to persist in the long term, irrespective of the process of degeneration. Last, an unspecified and unrelated selective advantage could be associated with recombination suppression. This would not be a mechanistic constraint, but we mention it as a possibility. It could for instance be the case for the maintenance of achiasmy in the heterogametic sex, independently of the evolution of sex chromosomes.

In this paper, we revisit this constraint scenario. We focus on the case where the short-term recombination arrest is caused by lucky inversions. The initial arrest is not the factor of interest here, so we use the simplest model (the lucky inversion process only requires the occurrence of deleterious mutations and variation in recombination rates). We contend that explanations based on the mechanistic constraint that recombination cannot be restored on the Y chromosome face several theoretical challenges, rendering them unlikely, in our view, to account for the evolution of sex chromosomes. Furthermore, we argue that mechanistic constraints on recombination restoration may often not be sufficiently strong to lead to stable heteromorphic sex chromosomes.

## Methods

We analyze a model of sex chromosome evolution where recombination arrest is caused by lucky inversions and explore the constraint scenario by varying the rate of recombination restoration. Specifically, we use the general model of sex chromosome evolution that we previously introduced to explore the regulatory theory [19,20, which should be consulted for more details], but removing the regulatory effects. This model considers a sex chromosome pair with a large number of genes (here 500) subject to deleterious mutations occurring in their coding sequences. Fitness is determined multiplicatively across loci by the effect of deleterious mutations with a dominance coefficient equal to 0.25, as observed on average for mildly deleterious mutations [40]. For simplicity, the SD locus is located at one extremity of the chromosome. Recombination variation is modeled by introducing mutations suppressing recombination in a region around the SD locus. These mutations can be thought of as being inversions (and we will refer to them as such) although other types of mechanisms are possible, as already mentioned. For instance, the removal of recombination hotspots, or the addition of a recombination suppressor sequence would work too. Specifically, as described in [17], we assume that inversions occur on the Y at a rate *U*_*inv*_ per chromosome per generation (we use *U*_*inv*_ = 10^−5^). We only consider inversions that include the SD locus (or extend the non-recombining region of the Y carrying the SD locus). Other inversions are not confined to males and can be fixed in the population, which does not lead to recombination suppression on the Y (homozygous inversions recombine normally). We denote the non-recombining fraction of the Y by *z* (between 0 and 1). This variable is also used to measure the endpoint of each inversion on the chromosome. When *z* = 0, X and Y chromosomes recombine freely, but otherwise X-Y recombination only occurs within the chromosomal segment [*z*, 1] (the SD locus being located at position 0). When *z* = 1, the X and Y do not recombine at all. When a new inversion occurs, its size is drawn as a uniform fraction of the non-recombining part of the Y. Specifically, on a Y where recombination is already stopped between 0 and *z_i_*, after a new inversion *i*+1 the non-recombining region will extend to *z*_*i*+1_ = *z*_*i*_ + (1 − *z*_*i*_)*u*, where *u* is a uniform deviate between 0 and 1. To allow for the possibility that recombination may be reestablished we assume that reversions can also occur (at a rate *U*_*rev*_ per chromosome per generation), reverting the last inversion on the non-recombining part of the Y. We investigate the dynamics of Y evolution in this model by supposing that reversion rates are much smaller than inversion rates, with *U*_*rev*_ = 10^−6^, 10^−7^, 10^−8^, 10^−9^, i.e., from 1 to 4 orders of magnitude lower than the rate of occurrence of inversions (*U*_*inv*_ = 10^−5^). At the start of a simulation, each individual carries a pair of fully recombining chromosomes, with the SD locus located at one extremity. Note that we do not perform a full exploration of the parameter space here, but rather use the simulations to illustrate the different points that are developed below concerning the effect of the reversion rate on sex chromosome evolutionary dynamics. Note that parameters scaled by the population size are likely to be the determinants of the evolutionary process so that the scenario can be extended to different population sizes by the appropriate rescaling of mutation rates, selection coefficients, and times. The simulations assume a constant population size (10^4^ offspring individuals are drawn each generation, irrespective of the average absolute fitness of male and female individuals in the previous generation). However, the population was considered to be extinct when the average fitness of males became a thousand times lower than the average fitness of females.

## Results

### Very low rates of reversion can prevent long-term recombination suppression

To evaluate how strong the constraint on reversions should be for recombination suppression to be maintained in the long term, we investigated cases where the rate of reversion was much lower than the rate of inversion. We ran replicated simulations lasting four million generations. With our standard parameters (Table 1), approximately 0.76 inversions are fixed per million generations. Typical outcomes are illustrated on Fig 1 (taken from runs with *U*_*rev*_ = 10^−8^). The majority of inversions that reach fixation remain relatively short-lived (Fig 2A) and most of them occur one at a time (i.e., they are reverted before a second one fixes). Before fixation, the marginal fitness of Y inversion decreases through time, as they tend to accumulate deleterious mutations (returning to the equilibrium load and being exposed to selective interference). After they fix, Y inversions continue degenerating because of selective interference, which reduces male fitness and eventually offsets their initial fitness advantage. At this point, they become deleterious, and it is just a matter of time before a reversion occurs that would be selectively favored (Fig 2B shows the marginal fitness at birth and at the time of reversion for fixed inversions, for different reversion rates). This is the example illustrated in Fig 1A. In a few cases, another inversion fixes on top of the first one, before the first has reversed (this corresponds to the example illustrated in Fig 1B). When the rate of reversion is very low, several inversions may stack on the first one. In these cases, the lifespan of the first inversion is prolonged because it can be reversed only after the second one (or third one, etc.) is reversed. This is due to the rather stringent hypothesis of our model that reversions can only occur on the last stratum present on the Y. This assumption protects the first inversion from reversion (until all other strata have reverted), and considerably reduces effective reversion rates (as it becomes zero for all but the last stratum on a Y). Considering reversions that could fully reestablish recombination on the Y at once would greatly reduce this (potentially unrealistic) effect. Whether or not stacking occurs, however, all inversions become reversed at some point (or the population goes extinct as we discuss below). There is no stable long-term maintenance of recombination suppression. Typically, Y chromosomes transiently carry one or two strata (Fig 4) for a relatively short time (Fig 2A). We can therefore conclude that the constraint scenario cannot explain the long-term maintenance of recombination suppression for rates of reversion up to 4 orders of magnitude lower than rates of inversion. This conclusion is very conservative, as our model of reversion does not allow the reversion of a first inversion if a second inversion occurs extending the non-recombining region (the first one can only be reversed after the second one is reversed). Without this constraint, it would be even more difficult to maintain recombination suppression.

**Fig 1.**
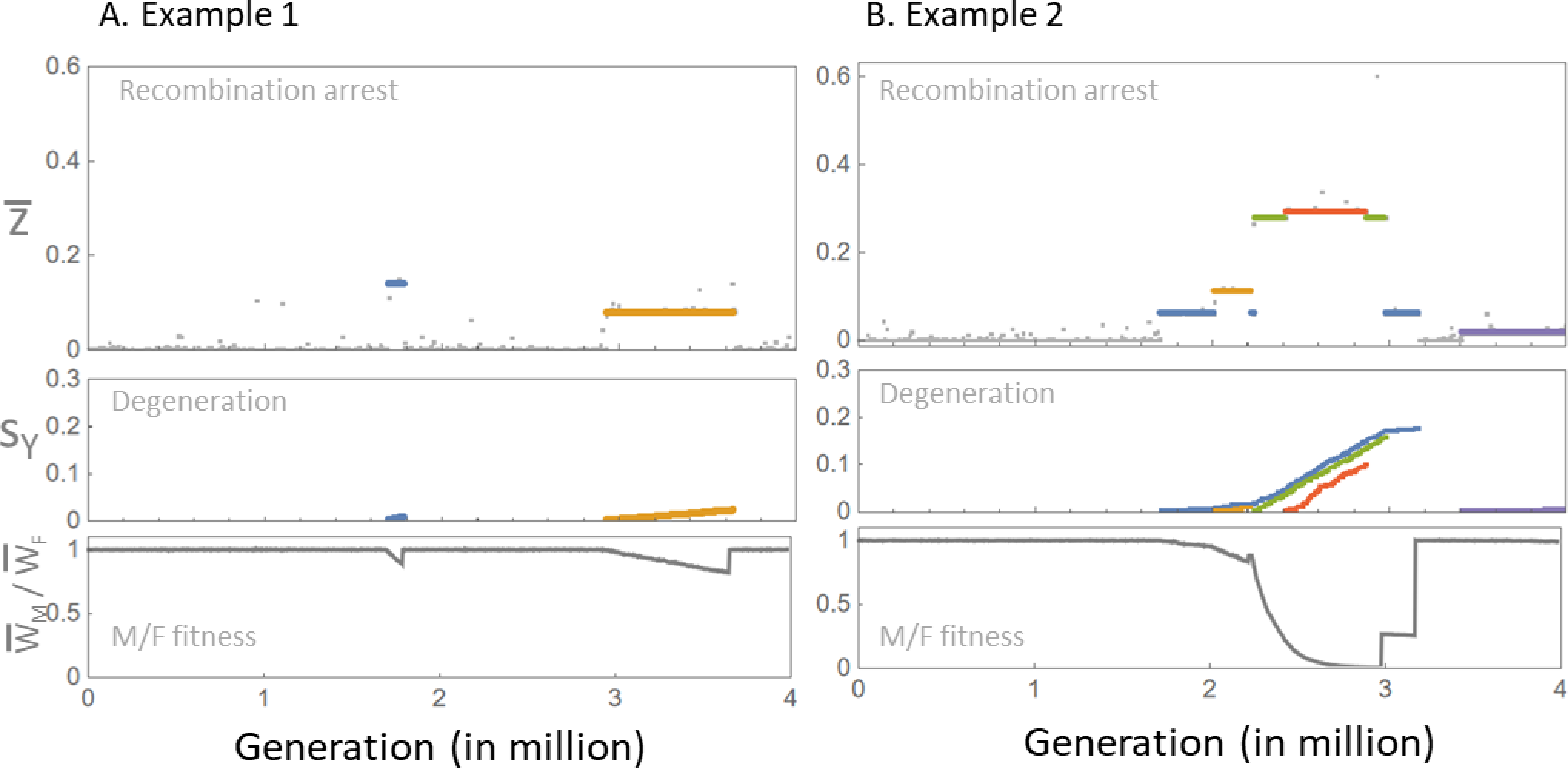
Examples of inversion-reversion dynamics. Each example is illustrated with three panels. The top panel shows the average nonrecombining fraction of the Y in the population (*z*) through time (x-axis in million generations). Colored lines correspond to fixed inversions (i.e., all inversions reaching a frequency of 1 during the simulation); different fixed inversions have different colors. The colored line extends between the time of occurrence of the inversion and the time when it becomes extinct. The middle panel shows the (per gene) average cumulative fitness effect of deleterious mutations on these inversions through time (same color code as in the top panel). The bottom panel shows the ratio of the average fitness of males/females in the population through time.

**Fig 2.**
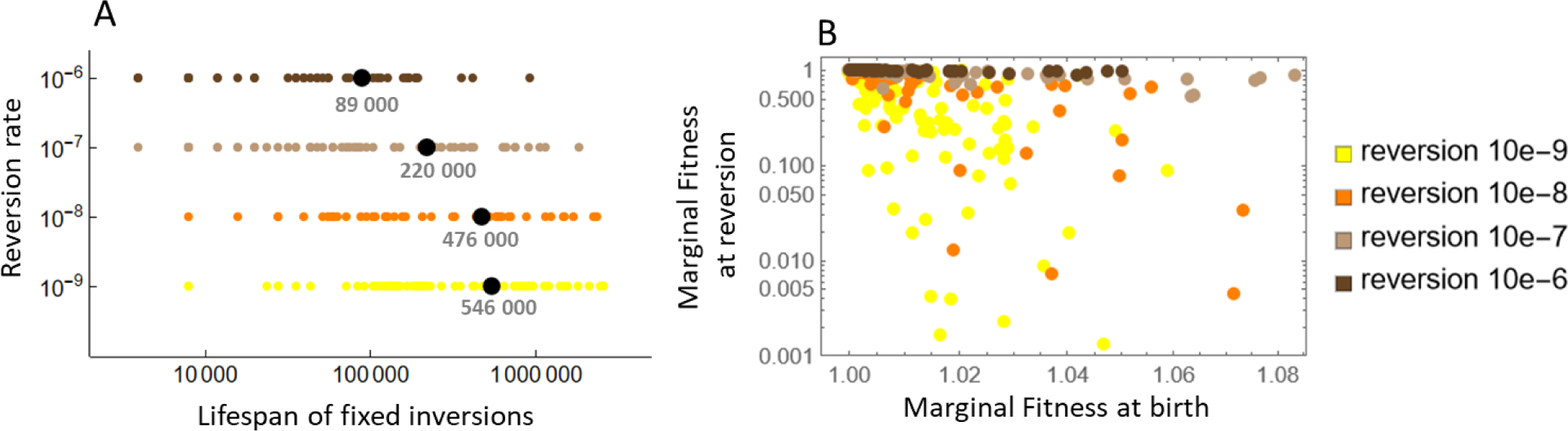
Characteristics of inversions under the constraint scenario. Panel A indicates the lifespan of fixed inversions (taken on 20 replicates) for different reversion rates (on the y-axis). Time was cut off at the end of the simulation (after 4 million generations) or if the population became extinct (by reaching a male/female fitness ratio < 0.001). Panel B shows the marginal fitness at birth of inversions (x-axis) versus marginal fitness at the last recorded time (y-axis, log scale). The latter most often corresponds to a reversion, but in a few cases, it corresponds to the end of the simulation or population extinction. Color codes correspond to different reversion rates as indicated in the legend.

**Fig 3.**
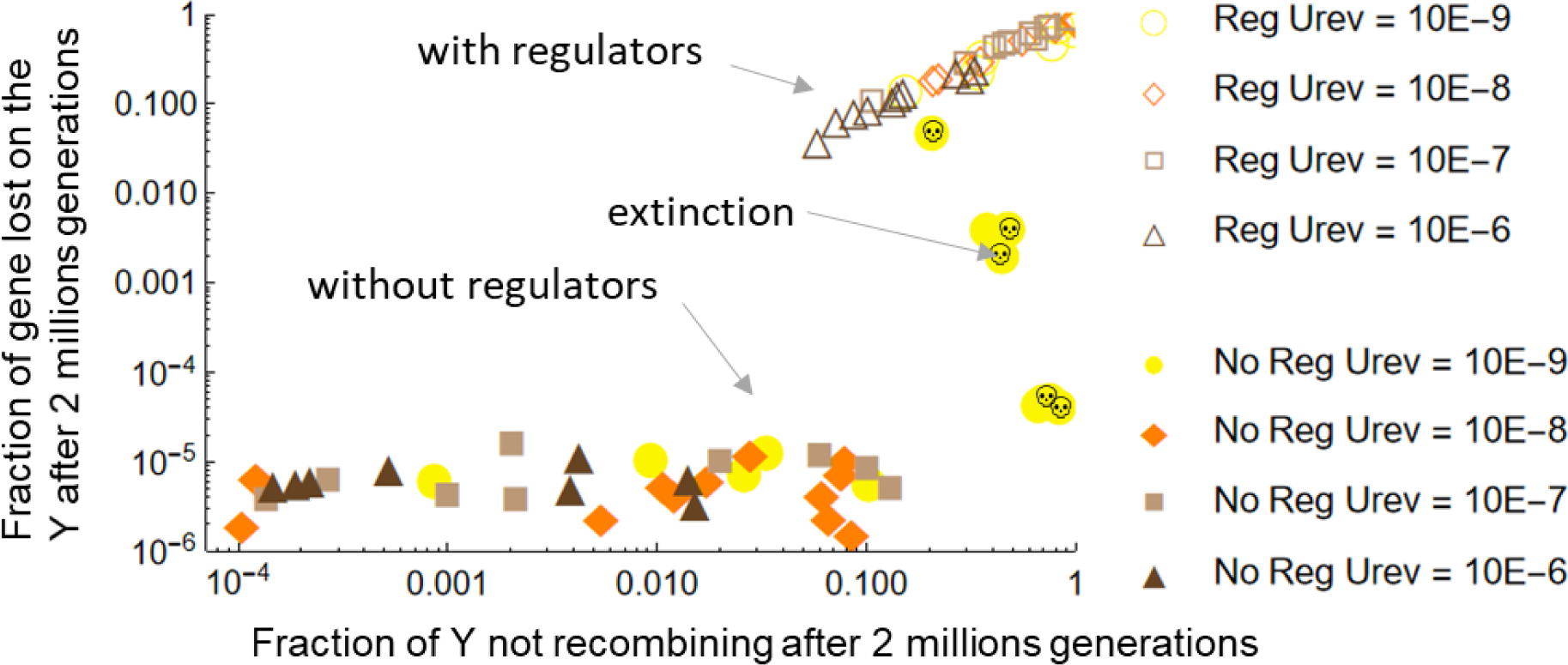
Overall evolution of the Y chromosome under the constraint scenario. The x-axis (in log scale) gives the fraction of the Y that is nonrecombining (averaged over all males in the population) after two million generations. The y-axis (in log scale) gives the fraction of genes lost on the Y after two million generations (averaged over all males in the population). A loss is defined as a gene having accumulated deleterious mutations up to *s_max_* = 0.3. Each dot represents a replicated population. Open symbols: regulators evolve (regulatory scenario); filled symbol: regulators do not evolve (constraint scenario). Color codes indicate different rates of reversion (*U_rev_*). The rate of inversion is 10^-5^ in all cases. In a few cases (with *U_rev_* = 10^-9^ in the constraint scenario, filled yellow disks) the population became extinct before 2 million generations. In these cases (marked with a skull), the x and y axes values are taken at the time of extinction.

**Fig 4.**
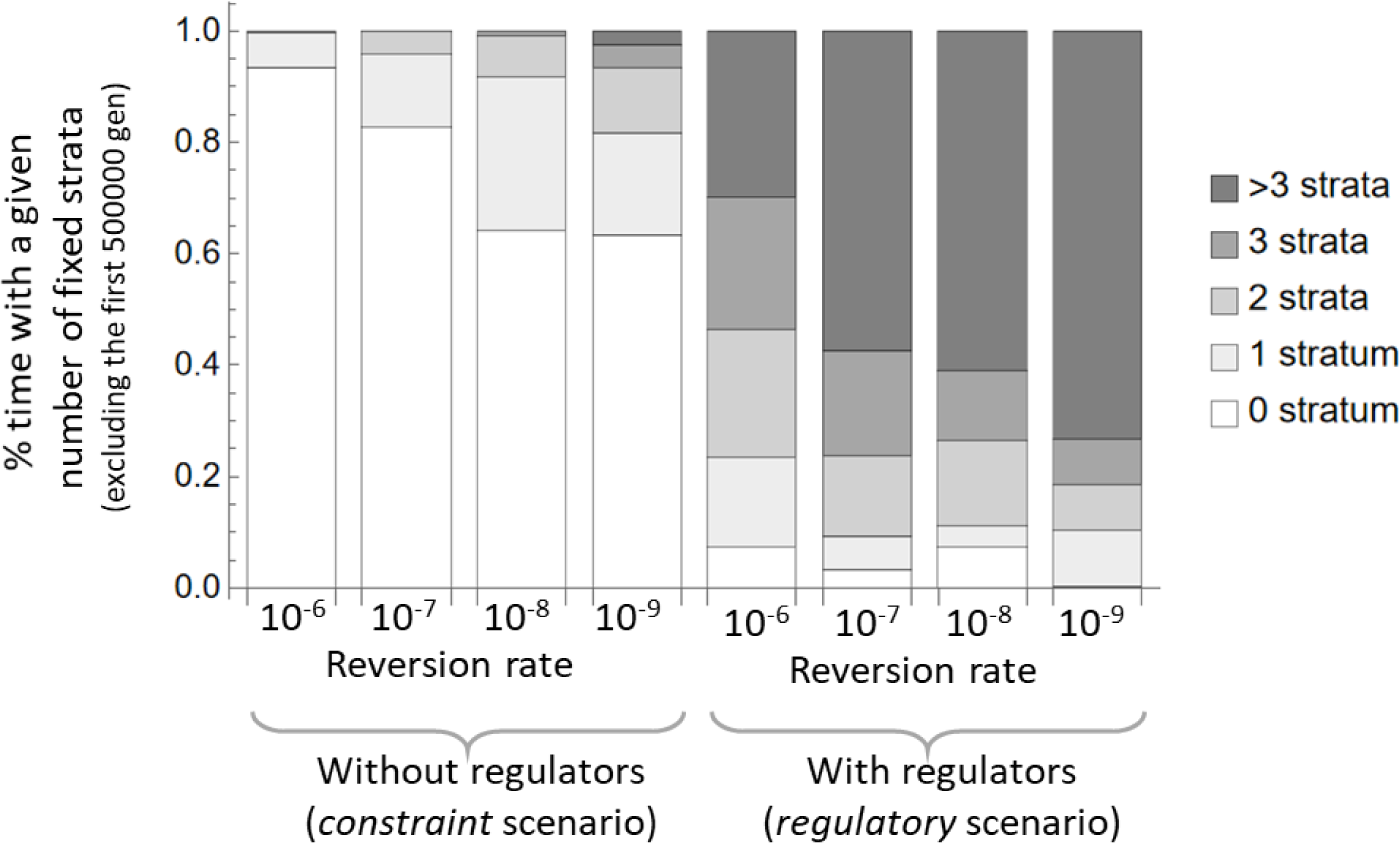
Fraction of the time during which the Y carries a given number of fixed strata. During a replicate simulation, there are times during which an inversion is fixed, perhaps including several strata, and times during which no fixed inversion is present (see examples in Fig 1). The bar chart gives this %time, across all replicates when a given number of strata are fixed in the population, for different reversion rates (without regulator evolution as in the constraint scenario, four bars on the left, or with regulator evolution as in the regulatory scenario, four bars on the right). This % is computed excluding the first 500 000 generation (to cut the initial phase influenced by the initial condition where the chromosome starts fully recombining and without any fixed inversion). The gray level corresponds to the number of fixed strata present as indicated in the legend on the right. For instance, in the constraint scenario with a reversion rate equal to 10^-6^, there is no fixed inversion 93.5% of the time, and one fixed inversion 6.3% of the time. In contrast, in the regulatory scenario with the same reversion rate, there is no fixed inversion 7.3% of the time, one fixed inversion 16% of the time, 2 fixed inversions 23% of the time, 3 fixed inversions 23.7% of the time and more than 3 fixed inversions 30% of the time (over the 3.5 million last generations of a simulation).

**Table 1.** Parameter values used in simulations. In grey, parameters are only used when regulators can evolve (in the constraint theory mutation rates on regulators are all set to zero).

| Parameters | Values |
| --- | --- |
| Population size | 10000 |
| Number of genes | 500 |
| Average effect of deleterious mutations (the distribution is exponential) | 0.05 |
| Dominance coefficient of deleterious mutations | 0.25 |
| Maximum fitness drop caused by mutations in a gene, corresponding to a gene loss ($s_{max}$) | 0.3 |
| Mutation rate of genes | 0.0002 |
| Recombination rate between consecutive genes ($R_g$) | 0.0005 |
| Rate of inversion mutations | 0.00001 |
| Rate of reversion mutations (10 to $10^4$ lower than the rate of inversions) | variable |
| Standard deviation of mutational effects on cis and trans-regulatory traits | 0.2 |
| Number of gene, cis-regulator, male-limited, and female-limited trans-regulators | 500 |
| Intensity of stabilizing selection on expression levels | 0.1 |
| Mutation rates on cis-regulators | 0.0002 |
| Mutation rates on trans-regulators | 0.0001 |
| Recombination rate between cis-regulator and its gene ($R_c$) | 0.00005 |

### Very low rates of reversion can prevent degeneration

Many old Y chromosomes are largely non-recombining and degenerate, mutations having accumulated up to the point where genes have become nonfunctional or have been lost. In our model, degeneration corresponds to the situation where a gene has accumulated deleterious mutations up to a maximum fitness effect of *s_max_* (corresponding to the fitness drop caused by the loss of function of a gene, here set to 0.3). When reversion rates are low, some inversions can fix and persist in the population for some time, especially when secondary inversions also occur prolonging the lifespan of the first one. Usually, this does not correspond to a large fraction of the Y, but on rare occasions, this can be significant (in particular, when several fixed inversions are stacked, which reduces the rate of reversion of all but the last one of them, as explained above). For example, in Fig 1B, about a third of the Y stopped recombining for nearly a million generations. Even in these extreme cases, however, degeneration remains moderate: even with extremely low rates of reversion (10^-9^), almost no gene accumulates deleterious mutations up to *s_max_*. Yet, we use a relatively high rate of deleterious mutation per gene (*U_g_* = 2 x 10^-4^), a distribution of fitness effects of mutations with a relatively high mean (*s_mean_* = 0.05), and a large proportion of small effect mutations (the distribution of effects is exponential). In the vast majority of cases, almost no loss of function is detectable (Fig 5A). The reason for this lack of loss of function is that many weakly deleterious mutations accumulate in all genes present on a fixed inversion. Collectively, their impact on the marginal fitness of the inversion starts to be strong long before any gene in particular becomes fully nonfunctional. Hence, inversions become selectively disfavored (and therefore selectively eliminated as soon as a reversion arises), long before they exhibit any gene loss. We can therefore conclude that the constraint scenario cannot explain strong degeneration for rates of reversion up to 4 orders of magnitude lower than rates of inversion. Degeneration may occur under very low rates of reversion if carrying nonfunctional genes on the Y would only cause a very small fitness cost for males (a situation that would be represented by setting *s_max_* to a small value in our model). This situation seems unlikely in the absence of a mechanism silencing impaired genes on the Y, however, while letting gene expression evolve would lead to the regulatory scenario, under which recombination arrest can be maintained even in the absence of any constraint on recombination restoration [17].

**Fig 5.**
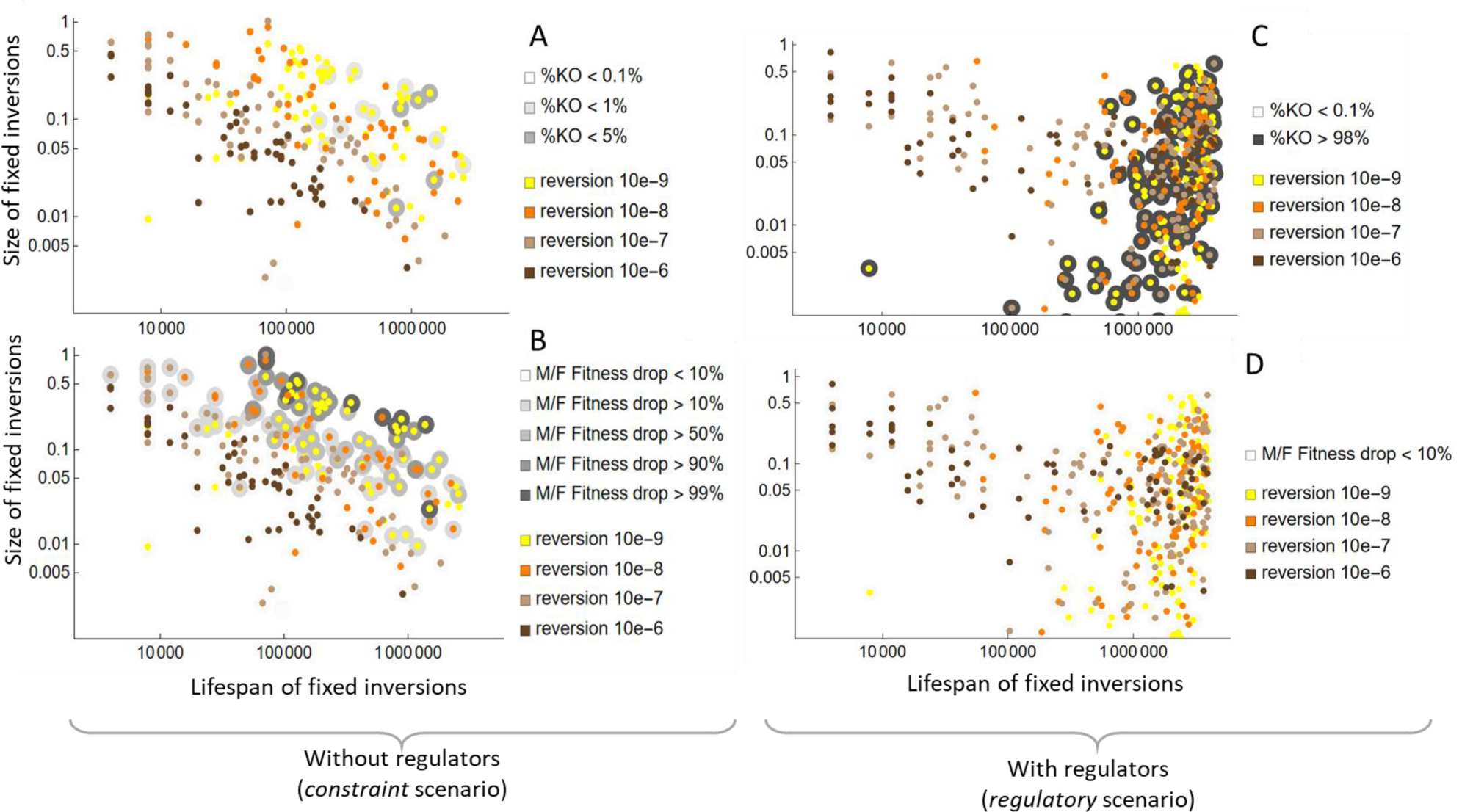
Detailed characteristics of fixed inversions in the constraint (panels A, B) and regulatory (panels C, D) scenarios. In all panels, the x-axis gives the lifespan of inversions (as defined in Fig 2A) and the y-axis the size of the inversion (both in log scale). Each dot represents a different fixed inversion that occurred across all replicates. The color of the dot indicates the reversion rate of the simulation during which the inversion was observed (as given in the legend). On panels A and C, a gray disk is added around each inversion. The gray level indicates the % of gene lost (%KO) on that inversion at the last time when the inversion is observed (i.e., just before it is reversed, at the end of the simulation at 4 million generations, or at population extinction). On panel A (constraint scenario), the gray level is light as this %KO never exceeds 5%. On panel C (regulatory scenario), this gray level is darker as this %KO reaches very high values (being either close to zero or above 98%). On panels B and D, a gray disk is also added around each inversion, this time representing the drop in the male/female fitness ratio caused by this inversion between the first and last time it is observed. Noting *r*(t) this male / female ratio at time *t*, this drop is computed as *r*(*t*_1_)/*r*(*t*_2_) between times *t*_1_ and *t*_2_. When several inversions are simultaneously present in a given time interval, the log(drop) is portioned proportionally to the relative size of each inversion *s*_1_/(*s*_1_+*s*_2_), i.e., with two inversions of size *s*_1_ and *s*_2_, the drop accrued to the first inversion is Exp( *s*_1_/(*s*_1_+*s*_2_) Log *r*(*t*_1_)/*r*(*t*_2_) ). For instance, with two inversions of equal size, each is assigned the square root of the fitness drop on the interval (so that the product of the fitness drop of each inversion gives the overall fitness drop). With this correction, the fitness drop associated with an inversion is more representative of what is happening on this inversion (rather than being caused by the presence of another inversion). On panel B (constraint scenario), these fitness drops can reach large values (more than a 99% reduction in the male/female fitness ratio), but they remain very low in the regulatory scenario (never exceeding 10% on panel D).

### The constraint scenario is more likely to lead to extinction than Y degeneration

It may be argued that reversion rates are even smaller than the ones we considered, making the constraint scenario a possibility, at least theoretically. There is a strong argument against this possibility. When an inversion fixes and starts accumulating deleterious mutations, it depresses male fitness (again, we take the example of XX/XY species, but the argument applies to the heterogametic sex: in ZZ/ZW species, females would show this fitness reduction). Initially, a lucky inversion is selectively favored because it captures a fraction of the Y carrying fewer deleterious mutations (or deleterious mutations with smaller effects) compared to the average Y population. Deleterious mutations start accumulating within the inversion because the inversion tends to return towards the average mutation load [18], and to selective interference. As explained above, a first threshold is reached when the accumulation of deleterious mutations depresses the marginal fitness of this portion of the Y below the average marginal fitness of the homologous portion of the X in the population. Reversions then become selectively favored, and it is just a matter of time before one occurs and eliminates the inversion. If the reversion rate is extremely low, this can indeed take a long time. However, a second threshold will be reached relatively quickly, corresponding to the non-viability or sterility of males, and hence to the extinction of the population/species. This cannot happen in our model as we assume a constant population size, i.e., the absolute number of individuals in the population does not depend on the fitness of individuals (soft selection), but the simulations show a crash in male fitness relative to female fitness when inversions are maintained for a sufficiently long time (Fig 5B). It is not easy to determine the male fitness threshold that would lead to population extinction in nature. We used a relatively conservative threshold equal to 10^-3^ (meaning that the fitness of males is three orders of magnitude lower than the fitness of females). Such a distortion of male vs. female fitness would be particularly conspicuous in natural conditions. In the few cases where some degeneration occurs (e.g., when a large inversion unfortunately fixes), the population reaches this limit quickly and becomes extinct. Reversions can rescue the population and prevent extinction, but if they are too rare, they do not occur quickly enough to prevent it. For instance, with very low rates of reversions (*U_rev_* = 10^-9^), this threshold was often reached in our simulations (extinction occurred in 70% of cases, 14 replicates out of 20 within the first 4 million generations of evolution). This estimate is conservative, since considering that the number of males in the population may be much lower than assumed under our soft selection regime would lead to an even faster accumulation of deleterious mutations on the Y (due to stronger drift). Note that our model does not include back mutations, which would eventually stop the decline in fitness caused by deleterious mutation accumulation. However, previous work has shown that in the absence of recombination, mean fitness reaches very low values even when back mutations do occur, unless the mean fitness effect of deleterious mutations is extremely weak [41,42]. Hence, a theory based on constraints alone cannot explain both degeneration and the persistence of populations/species. For degeneration and persistence to occur, Y silencing and DC must also evolve, which can be a powerful selective mechanism that stabilizes recombination arrest.

### Comparing the constraint scenario to a scenario including a selective pressure against recombination

It is not because reversion rates are low (representing strong constraints on recombination restoration) that explanations of long-term recombination suppression solely based on constraints are likely to hold. Quantitatively, the question is rather, for given reversion rates, to determine the most likely scenario for observing Y chromosomes with non-recombining and degenerate strata within a realistic timeframe. To illustrate this point, we simulated the evolution of Y chromosomes under the same low reversion rates used above, but allowing regulators to evolve using the model described in [17], which should be consulted for more details. We use the same simulations as above, but we consider that the expression of each gene is controlled by a cis-regulator and two trans-regulators (one only expressed in males and the other only expressed in females). These regulators determine quantitative traits that control the total and allele-specific level of expression of each gene. Total gene expression is supposed to be under stabilizing selection for all genes. Fitness is determined multiplicatively across loci by the effect of deleterious mutations (whose dominance depends on the relative strength of cis-regulators in heterozygotes, with a baseline dominance in the absence of cis-regulatory variation equal to 0.25) and by the departure from optimal expression at each locus. Cis and trans regulators mutate at a fixed rate per generation (with Gaussian variation in trait values). Table 1 indicates the additional parameters and their value for the simulations with the evolution of regulators.

In this case, we observe much faster Y recombination suppression and degeneration than in the absence of regulatory evolution, for all reversion rates investigated. This is probably also true for scenarios involving SA loci, which generate a selective pressure against recombination accelerating the process. Low reversion rates are favorable to any theory on the maintenance of recombination arrest, not only those solely based on constraints. Indeed, low reversion rates give more time for other SA loci to accumulate, or nascent DC to emerge, and more time for degeneration to happen. In our model of regulatory evolution, and with very low reversion rates, almost any fixed inversion has time to develop nascent DC, generating sexually antagonistic regulatory effects that effectively disfavor recombination. Fig 4 shows that many more strata accumulate on the Y in this case than in the absence of regulatory evolution, while Fig 5C shows that most of these strata are long-lived and fully degenerated (while almost none is degenerated for the same parameter values in the absence of regulatory evolution, Fig 5A). Fig 5D shows that this degeneration is not associated with a large drop in male-to-female fitness ratio, while this drop is considerable in the absence of regulatory evolution. When regulators evolve, reversions occur but are not selectively favored. Indeed, the reestablishment of X-Y recombination on a given stratum causes X cis-regulators to move to the Y, creating recombinant (low fitness) Y that cause a departure from optimal gene expression in males. This is a case of an evolved selective constraint. Strata are selectively stabilized because DC emergence creates sex antagonistic regulatory effects on expression levels, as well as silencing of the deleterious mutations accumulating on the Y. Hence, strata become permanently stabilized and can persist indefinitely, in contrast to the constraint theory where strata are never stable. The ultimate cause of long-term recombination suppression is not the absence of genetic variation for reestablishing recombination once an inversion has fixed (mechanistic constraint), but that it is selectively unfavorable to reestablish it. Overall, for the parameter values considered here, we see that after 2 million generations a large fraction of the Y has stopped recombining and degenerated in the presence of regulatory evolution with the lowest reversion rates (for high reversion rates the process is still ongoing with only a few small stabilized and degenerated strata, Fig 3). Nothing nearly comparable occurs without regulatory evolution, where at most, in a few cases with extremely low rates of reversion, some recombination suppression evolves and drives the population to extinction without leading to significant Y degeneration (Fig 3). We conclude that even if reversion rates were extremely low so that a scenario solely based on constraints could produce partial and transient degeneration and recombination suppression, a scenario involving a selective pressure maintaining recombination arrest is orders of magnitude more likely to produce complete and permanent recombination suppression and degeneration, without a major drop in male fitness.

## Discussion

The arguments and results presented in this article imply that, in the absence of regulatory evolution, the decreased fitness of the heterogametic sex due to mutation accumulation on the Y should lead to two possible outcomes: (i) the restoration of recombination if reversions can occur, even at very low rates, or (ii) the extinction of populations if constraints on reversions are sufficiently strong. In the following, we discuss the constraint theory in the light of those results and possible mechanisms of recombination reestablishment, before indicating future avenues for theoretical and empirical research concerning the initial steps of recombination suppression, the mechanisms responsible for the long-term maintenance of recombination suppression, and the extension of these theories.

### What is the degree of constraint on recombination reestablishment?

The possibility of recombination restoration should depend on the mechanism of recombination arrest. Recent theoretical work has emphasized the possible role of inversions in suppressing recombination on sex chromosomes [17–19,43], although these models also apply to other mechanisms of stepwise recombination arrest. Inversions do indeed occur frequently within populations and may be caused by ectopic recombination between repeated sequences [44–47]. For this reason, they often tend to occur on the same sites and sometimes repeatedly [48–50]. They are often observed on sex chromosomes [10,51–53], although some of these inversions may have occurred after recombination arrest [10]. Inversions are well known to reduce recombination rates in heterokaryotes. This reduction is not necessarily because inversions inhibit homologous pairing. If inversions are sufficiently large, pairing can occur, and inversions form loops allowing for a local alignment of the two homologous chromosomes. These loops can directly inhibit chiasma formation, especially near the breakpoints of the inversion [54,55, but see 56]. However, the suppression of recombination is also strongly mediated by the fact that an odd number of crossovers within the inversion loop leads to the production of unbalanced chromosomes. Such unbalanced chromosomes usually cause a fitness reduction and are thus eliminated (however, as explained in the appendix, this may be less true when the chromosome is degenerated). Finally, it is important to note that recombination may still occur when the number of crossovers falling within the inversion is even (especially in the case of relatively large inversions where crossover interference is weaker [57–60]), while gene conversion events may also allow for genetic exchanges between inverted and non-inverted segments [54,60]. Such exchanges would limit degeneration and thus allow for longer persistence time of inversions compared to the situation modeled in our simulations.

In a previous model of inversion dynamics on sex chromosomes, Jay et al. [19] consider that recombination restoration is possible only when an inversion with exactly the same breakpoints (“reinversion”) restores the exact original gene order before another inversion overlaps or occurs within the first one. If a second nested or overlapping inversion occurs, it can also be reversed, but only before a third nested/overlapping inversion occurs, and so on. In this model, the chance of reinversion becomes vanishing low as the number of breakpoints increases. Indeed, with *N* breakpoints on the chromosome and with a first inversion spanning *k* breakpoints, the chance of reinversion is ∼1/*N*^2^, while the chance of a second overlapping / nested inversion is ∼*k*/*N*. In the results we presented, this level of constraint is achieved for the very low rates of reversions. For instance, with *N* = 100 breakpoints and a first inversion spanning *k* = 10 breakpoints, reinversions are 3 orders of magnitude less likely than inversions. An interesting feature of this model is that it mechanistically represents inversions and reinversions. It also captures the idea that reestablishing the exact gene order with random inversions becomes increasingly difficult as they accumulate. This phenomenon may indeed occur and constrain the reestablishment of recombination.

However, several processes could largely limit the constraint imposed by the accumulation of overlapping and nested inversions. First, recombination may occasionally occur even if gene collinearity is not exact, as shown with ectopic recombination [61–63], especially after a second inversion restoring the original direction on a portion of the chromosome. Fig S1 and S2 in the appendix illustrate such possibilities for nested or overlapping inversions. With imperfect collinearity, recombined chromosomes have low fitness in general, but here, with a partially degenerated Y, the question is more subtle, as the loss of some (already) degenerated genes may be compensated by the acquisition of non-degenerated portion of the X. What matters is the relative fitness of recombined Y compared to the current (partially degenerated) low fitness Y. Hence, many favorable cases of imperfect recombination could occur and be favored, which could largely increase rates of recombination reestablishment.

Another possibility is that recombining sex chromosomes may be reestablished by moving the SD locus out of the non-recombining region. This mechanism can always occur, even when complex rearrangements have taken place on the Y. This may occur for instance by recruiting a new master switch gene for sex-determination or following a duplication of the existing SD locus into a new location. Gene duplications are frequent events in eukaryotic genomes [64]. Rates in the range 10^-5^ – 10^-7^ per gene per generation have been reported in animals [65–67], i.e. a much higher rate than the rate of reversions that we investigated. If the SD locus moves to another (recombining) location on the sex chromosome, a new Y recently re-derived from the X (and fully recombining) can evolve easily. Examples of such events are often reported [68–70], indicating that they may be common. The SD locus may also move to another autosomal pair, leading to the evolution of a new pair of fully recombining sex chromosomes. Such turnovers of sex chromosomes have also been reported in several species [71,72] and are predicted to be favored when deleterious mutations have accumulated due to the lack of recombination [37,38]. In some species, another possibility for restoring recombination on the Y involves environmental sex reversal when recombination is sex-dependent [36,73,74].

Often, the idea that recombination restoration is strongly constrained stems from the idea that ‘reinversions’ are not observed. While the rate of reinversion is probably low, this view is not entirely accurate. Reinversions have been occasionally shown to occur in laboratory populations of Drosophila [75] at rates 10^-3^ – 10^-4^: while most of them were caused by X-ray irradiation [76,77], one occurred spontaneously in a stock population [78,79] and it was proposed that reinversion may be favored by the physical proximity of the breakpoints during loop formation [77]. Recent comparative genomics has also highlighted that inversions and reinversions occur frequently and repeatedly at particular breakpoints, although estimating the corresponding rates has not been done [49,50]. More empirical work on this issue is needed to assess to what extent such a process may occur (and at which rate) despite the repression of crossing-over in the vicinity of inversion breakpoints. However, the observation of an accumulation of nested and overlapping inversions alone is not an indication that the absence of recombination restoration was due to a constraint or to a selection pressure against recombination. Chromosomal rearrangements can secondarily accumulate in a non-recombining region [1,10].

Hence, while the accumulation of complex rearrangements is certainly a way to suppress recombination, the maintenance of recombination arrest on the Y by a constraint alone requires a very high level of constraint, given the maladaptation caused by mutation accumulation on the Y. This level of constraint is, in fact, a rather strong assumption. It requires that reinversions are very rare, that rare recombination events involving imperfectly colinear chromosomes do not occur, and that the SD locus cannot move outside the non-recombination region. In any case, determining whether recombination suppression is maintained selectively or by a mechanistic constraint is likely to be empirically difficult. A key difference between these two cases is that Y strata have a higher or lower marginal fitness compared to the equivalent segment on the X. If it was feasible to experimentally switch these portions of chromosome and investigate the fitness effect of this switch, the two cases would lead to opposite predictions. In the constraint theory, the switch should increase male fitness, while the opposite is expected if recombination suppression is selectively maintained, provided some degeneration has occurred.

### Conclusion and perspectives

After recombination suppression, the accumulation of deleterious mutations on the Y generates a selective pressure to restore recombination and purge those alleles. This selective pressure becomes stronger as male fitness declines, soon making recombination restoration events highly favorable. This explains that even with extremely low rates of recombination reestablishment, recombination suppression cannot persist in the long term when only deleterious mutations are considered. In this case, recombination restoration events are rescuing the population from extinction, and even if they are rare, play a disproportionate role in the outcome. This issue is even more acute in models where recombination arrest is caused by neutral divergence [22] as recombination suppression occurs gradually, rather than quickly as in the lucky inversion scenario. This is the central theoretical argument against theories of sex chromosome evolution solely based on mechanistic constraints [19,22]: In the absence of regulatory evolution, the accumulation of deleterious alleles caused by recombination arrest should eventually lead to population extinction or the reestablishment of recombination (via reinversion(s), or via a change of location of the sex-determining locus, either to the PAR or to an autosome as with a sex chromosome turnover), rather than the long-term maintenance of degenerate Y or W chromosomes.

Several issues remain to be investigated in more detail. The different processes possibly involved in the evolution of Y chromosomes need to be better integrated. In particular, the conditions allowing the long-term maintenance of recombination suppression under the SA theory should be investigated further [following 37,38] and combined with models of regulatory evolution. Better integration of the different mechanisms of regulatory variation may also be useful, such as mechanisms based on imprinting [80] or based on the reallocation of transcription factors to X-linked genes [11]. While the widespread occurrence of DC [80,81] indicates that it is needed at least for some genes, it would be interesting to introduce heterogeneity among genes in selection on dosage, and varying the genetic architecture of DC (local vs global). Investigating whether DC always evolves, at least for dosage-sensitive genes, for old degenerate sex chromosomes would also be interesting, to confirm the key role of DC in their long-term stability. Several cases of interest should be investigated further, notably cases involving non-random mating [18,29,82] or UV and mating-type chromosomes. Analytical models are needed to generalize the recent findings that have mostly been explored by simulation. Finally, from an empirical perspective, more data are needed on patterns of recombination suppression, degeneration, and mechanisms of early DC evolution in young sex chromosome systems. Overall, the level of constraint on recombination restoration may not be the key parameter to understand why heteromorphic or homomorphic sex chromosomes occur in a given species. The ease of evolving dosage compensation is likely to be the main driver: if regulatory evolution is difficult, sex chromosomes will remain homomorphic (recombination will be restored in some way or the species will go extinct). If regulatory evolution and DC can evolve relatively easily, stable heteromorphic sex chromosomes may persist in the long term [17].

## Acknowledgments

We thank A. Muyle, J. Käfer and anonymous reviewers for useful comments and suggestions. We thank L. Ferretti, M. Kirkpatrick, and S. Otto for useful discussions.

## Funding

This work is supported by the ANR Genasex, the ANR CisTransEvol and the ERC RegEvol.

### Conflict of interest disclosure

The authors declare that they comply with the PCI rule of having no financial conflicts of interest in relation to the content of the article. TL and DR are recommenders for PCI Evol. Biol.

### Code availability

The simulation code is available at:

T. Lenormand, D. Roze, Zenodo (2021), doi:10.5281/zenodo.5504423.

## Appendix

### Recombination reestablishment after secondary nested or overlapping inversions

A difficulty to model recombination evolution by inversions and reinversions is that it is difficult to model the fact that the recombination process is not ‘perfect’ in the sense that it can occur between regions that are only locally homologous, as in the case of ectopic recombination. In the case of Y evolution, recombination may occasionally occur even in the absence of exact collinearity between the X and Y. This is illustrated in Fig S1 for an overlapping inversion that includes the SD locus, and in Fig S2 for nested inversions. In both cases, the secondary inversion has a homologous region on the X and can pair at meiosis, allowing recombination to occur around the SD locus. If a crossover occurs within this region, a new Y can be produced which may not carry the exact complement compared to the original chromosome (with either deleted or duplicated positions). In each case, such a crossover will eliminate parts of the former Y on the first stratum, i.e., regions that may have already accumulated a load of deleterious mutations. This recombined Y could be particularly favorable (compared to the degenerated Y), even if some sequences are duplicated or missing compared to the X (see appendix and Figs S1, S2). Indeed, whether the recombined Y can invade depends on its marginal fitness relative to the marginal fitness of the potentially highly degenerated Y chromosomes present in the population (and not to the marginal fitness of a hypothetical mutation-free Y chromosome with full gene content). Furthermore, the recombined Y may be “improved” in further steps since it can now recombine more easily with the X around the SD locus after this first recombination event. In particular, a second crossover near the SD locus can further improve collinearity with the X and eliminate further degenerated parts of the Y from the first stratum that are still present (see Figs S1, S2). Alternatively, recombination may also be restored if an inversion arises on the X, facing the inversion on the Y [17]. Again, more empirical work is needed to assess whether recombination may indeed occur in such scenarios despite non-perfect collinearity. However, the occurrence of ectopic recombination between repeated sequences indicates that it is possible in principle [45,62], while the results of the present article show that even very low rates would be sufficient to maintain recombination in the long term.

---

**Fig. S1.**
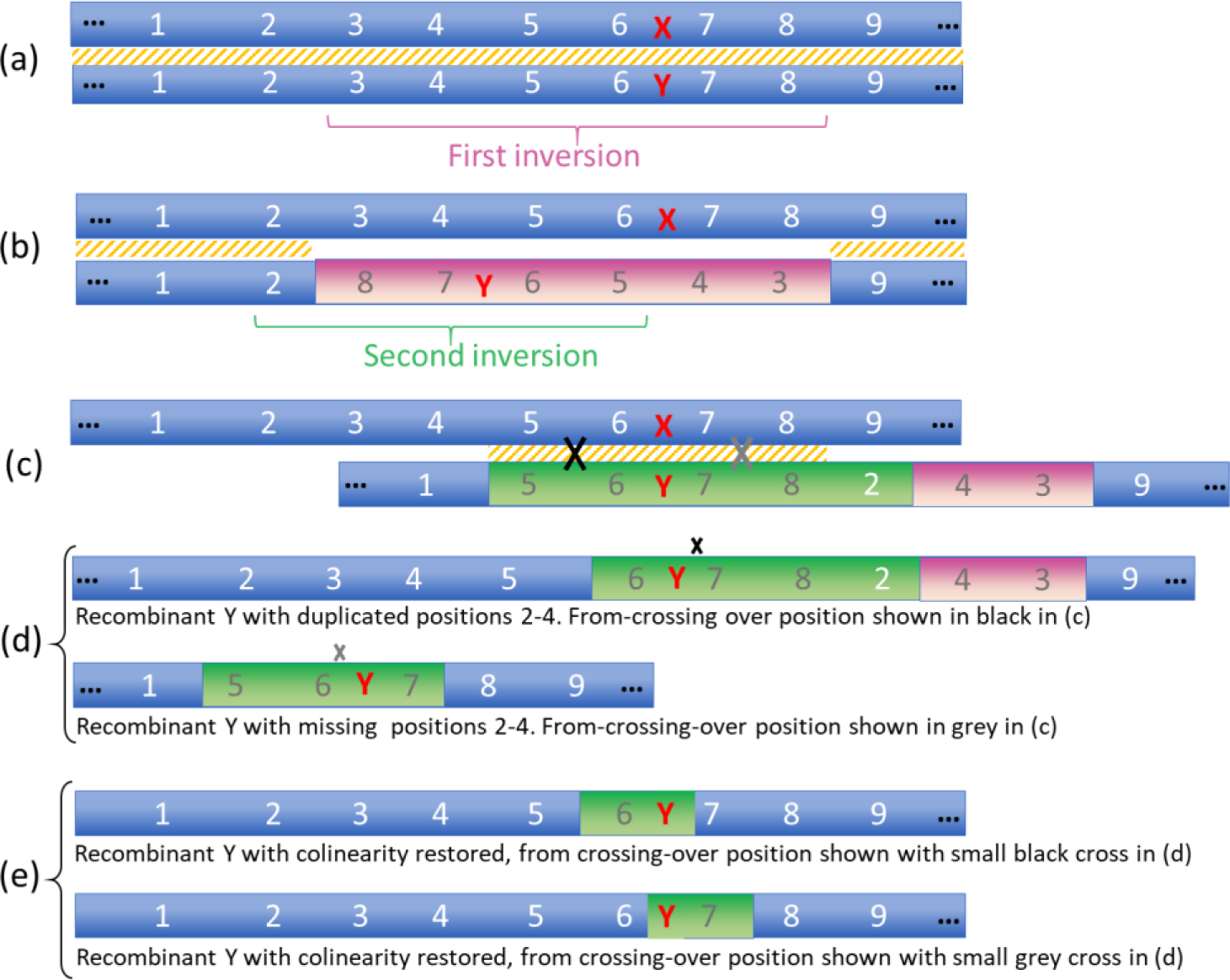
Restoration of recombination on the Y with overlapping inversions. (a) depicts the fully recombining XY pair with the SD locus in red. The orange dashing indicates where homologous pairing is possible. A first inversion occurs (purple) on the Y between positions 3 and 8 leading to the situation in (b). Pairing and recombination do not occur around the SD locus. This first stratum on the Y chromosome can start degenerating in the absence of recombination. This is shown by the brown color of the position numbers. A second overlapping inversion occurs (green) on the Y between positions 2 and 5. The resulting Y in (c) can now pair with the X between positions 5 and 8. Crossing-over on the left of the SD locus (black cross) can generate recombinant Y with duplicated positions 2-4 (note that this Y recovers functional copies in positions 3-5, and has two functional copies in position 2). Crossing-over on the right of the SD locus (gray cross) can generate recombinant Y with missing positions 2-4 (note that the lack of these positions may not reduce fitness a lot if they were degenerated). (e) In both cases, a second crossover with the X occurring near the SD locus (shown with a small cross) can restore full collinearity with the X and get rid of other degenerated copies (on the right and left position of the Y, respectively).

**Fig. S2.**
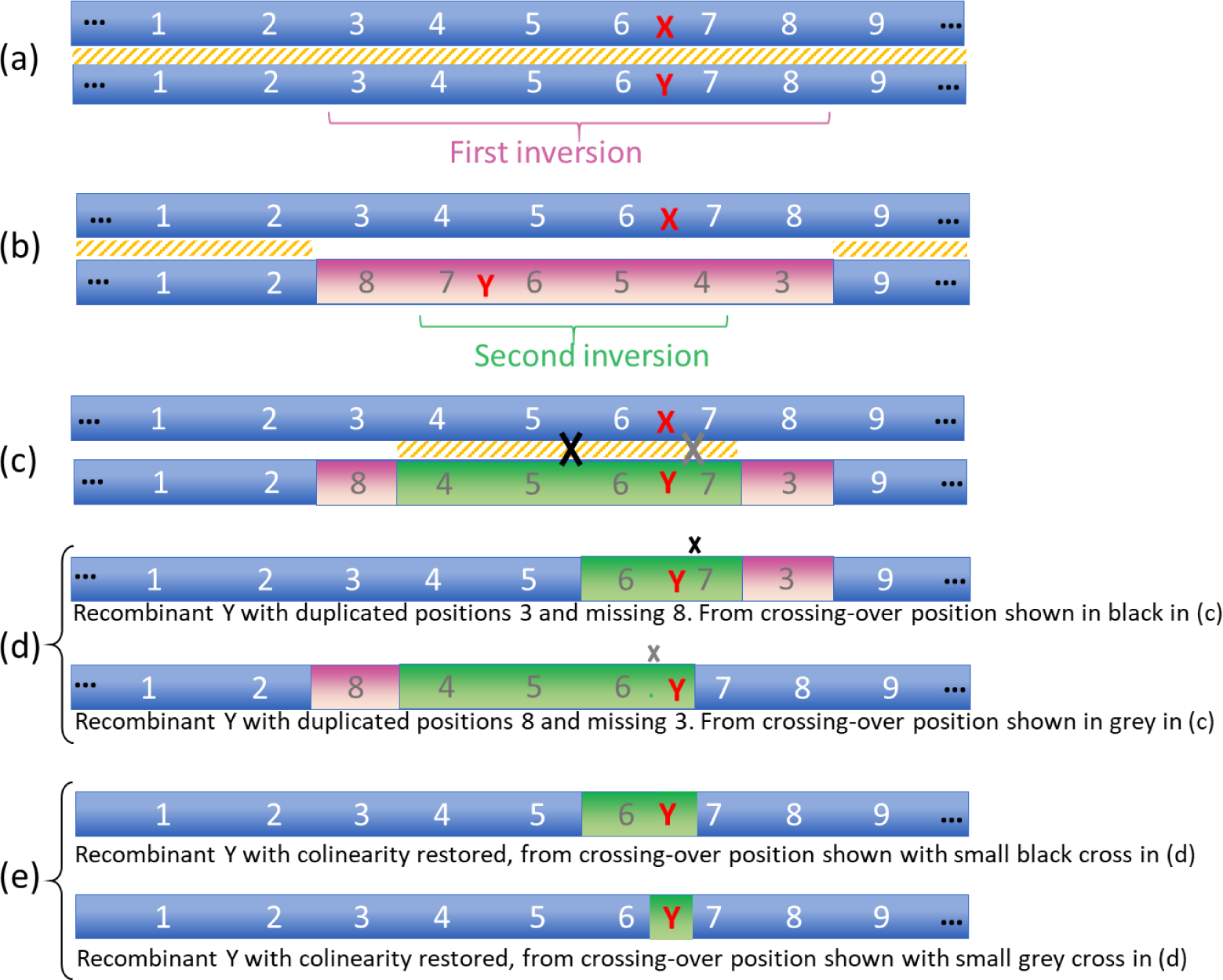
Restoration of recombination on the Y with nested inversions. (a) depicts the fully recombining XY pair with the SD locus in red. The orange dashing indicates where homologous pairing is possible. A first inversion occurs (purple) on the Y between positions 3 and 8 leading to the situation in (b). Pairing and recombination do not occur around the SD locus. This first stratum on the Y chromosome can start degenerating in the absence of recombination. This is shown by the brown color of the position numbers. A second nested inversion occurs (green) on the Y between positions 4 and 7. The resulting Y in (c) can now pair with the X between positions 4 and 7. Crossing-overs on the left of the SD locus (black cross) can generate a recombinant Y with duplicated positions 3 and missing 8. Note that this Y can have a higher fitness by recovering functional copies at positions 3-5. Position 8 is missing, but this may not be consequential since it was a degenerated position. Crossing-over on the right of the SD locus (gray cross) can generate a recombinant Y with duplicated position 8 and missing 3. Again, the fitness of this Y may increase, since it recovers functional copies at positions 7-8 while losing a degenerated copy at position 3. (e) In both cases, a second crossover with the X occurring near the sex-determining locus (shown with a small cross) can reconstitute a Y chromosome fully colinear with the X.

## Notes

### Competing Interest Statement

The authors have declared no competing interest.

### Summary of Updates

spelling errors

## References

1. Charlesworth D. When and how do sex-linked regions become sex chromosomes? Evolution. 2021;75: 569–581. doi:10.1111/evo.14196

2. Rifkin JL, Beaudry FEG, Humphries Z, Choudhury BI, Barrett SCH, Wright SI. Widespread recombination suppression facilitates plant sex chromosome evolution. Molecular Biology and Evolution. 2021;38: 1018–1030. doi:10.1093/molbev/msaa271

3. Bell G. The masterpiece of nature: the evolution and genetics of sexuality. Berkeley: University of California Press; 1982.

4. Lenormand T, Dutheil J. Recombination difference between sexes: a role for haploid selection. Hurst LD, editor. PLoS Biology. 2005;3: 396–403. doi:10.1371/journal.pbio.0030063

5. Charlesworth B, Charlesworth D. A model for the evolution of dioecy and gynodioecy. The American Naturalist. 1978;112: 975–997. doi:10.1086/283342

6. Bull JJ. Evolution of sex determining mechanisms. Menlo Park, CA: Benjamin Cummings; 1983.

7. Rice WR. Sex chromosomes and the evolution of sexual dimorphism. Evolution. 1984;38: 735– 742.

8. Rice W. The accumulation of sexually antagonistic genes as a selective agent promoting the evolution of reduced recombination between primitive sex chromosomes. Evolution. 1987;41: 911–914.

9. Charlesworth B. The evolution of sex-chromosomes. Science. 1991;251: 1030–1033.

10. Charlesworth D. Evolution of recombination rates between sex chromosomes. Phil Trans R Soc B. 2017;372: 20160456. doi:10.1098/rstb.2016.0456

11. Charlesworth B. Model for evolution of Y chromosomes and dosage compensation. Proceedings of the National Academy of Sciences of the United States of America. 1978;75: 5618–5622.

12. Bachtrog D. Y-chromosome evolution: Emerging insights into processes of Y-chromosome degeneration. Nature Reviews Genetics. 2013;14: 113–124. doi:10.1038/nrg3366

13. Lenormand T. The evolution of sex dimorphism in recombination. Genetics. 2003;163: 811–822.

14. Rowe L, Chenoweth SF, Agrawal AF. The genomics of sexual conflict. American Naturalist. 2018;192: 274–286. doi:10.1086/698198

15. Bonduriansky R, Chenoweth SF. Intralocus sexual conflict. Trends in Ecology and Evolution. 2009;24: 280–288. doi:10.1016/j.tree.2008.12.005

16. Ironside JEE. No amicable divorce? Challenging the notion that sexual antagonism drives sex chromosome evolution. BioEssays. 2010;32: 718–726. doi:10.1002/bies.200900124

17. Lenormand T, Roze D. Y recombination arrest and degeneration in the absence of sexual dimorphism. Science. 2022;375: 663–666. doi:10.1101/2021.05.18.444606

18. Olito C, Ponnikas S, Hansson B, Abbott JK. Consequences of partially recessive deleterious genetic variation for the evolution of inversions suppressing recombination between sex chromosomes. Evolution. 2022;76: 1320–1330. doi:10.1111/evo.14496

19. Jay P, Tezenas E, Véber A, Giraud T. Sheltering of deleterious mutations explains the stepwise extension of recombination suppression on sex chromosomes and other supergenes. PLOS Biology. 2022;20: e3001698. doi:10.1371/journal.pbio.3001698

20. Nei M, Kojima KI, Schaffer HE. Frequency changes of new inversions in populations under mutation-selection equilibria. Genetics. 1967;57: 741–750.

21. Bowen ST. The genetics of *Artemia salina*. V. Crosssing over between the X and Y chromosomes. Genetics. 1965;52: 695–710.

22. Jeffries DL, Gerchen JF, Scharmann M, Pannell JR. A neutral model for the loss of recombination on sex chromosomes. Philosophical Transactions of the Royal Society B: Biological Sciences. 2021;376. doi:10.1098/rstb.2020.0096

23. Datta A, Hendrix M, Lipsitch M, Jinks-Robertson S. Dual roles for DNA sequence identity and the mismatch repair system in the regulation of mitotic crossing-over in yeast. Proceedings of the National Academy of Sciences. 1997;94: 9757–9762. doi:10.1073/pnas.94.18.9757

24. Li L, Jean M, Belzile F. The impact of sequence divergence and DNA mismatch repair on homeologous recombination in *Arabidopsis*. The Plant Journal. 2006;45: 908–916. doi:10.1111/j.1365-313X.2006.02657.x

25. Do AT, Brooks JT, Le Neveu MK, LaRocque JR. Double-strand break repair assays determine pathway choice and structure of gene conversion events in *Drosophila melanogaster*. G3 Genes|Genomes|Genetics. 2014;4: 425–432. doi:10.1534/g3.113.010074

26. Benavente E, Sybenga J. The relation between pairing preference and chiasma frequency in tetrasomics of rye. Genome. 2004;47: 122–133. doi:10.1139/g03-134

27. Blackwell AR, Dluzewska J, Szymanska-Lejman M, Desjardins S, Tock AJ, Kbiri N, et al. MSH 2 shapes the meiotic crossover landscape in relation to interhomolog polymorphism in *Arabidopsis*. EMBO J. 2020;39. doi:10.15252/embj.2020104858

28. Ziolkowski PA, Berchowitz LE, Lambing C, Yelina NE, Zhao X, Kelly KA, et al. Juxtaposition of heterozygous and homozygous regions causes reciprocal crossover remodelling via interference during *Arabidopsis* meiosis. eLife. 2015;4: e03708. doi:10.7554/eLife.03708

29. Charlesworth B, Wall JD. Inbreeding, heterozygote advantage and the evolution of neo–X and neo–Y sex chromosomes. Proc R Soc Lond B. 1999;266: 51–56. doi:10.1098/rspb.1999.0603

30. Rice WR. Genetic hitchhiking and the evolution of reduced genetic activity of the Y sex chromosome. Genetics. 1987;116: 161–167. doi:10.1093/genetics/116.1.161

31. Peck JR. A ruby in the rubbish: beneficial mutations, deleterious mutations and the evolution of sex. Genetics. 1994;137: 597–606.

32. Charlesworth B. The evolution of chromosomal sex determination and dosage compensation. Current Biology. 1996;6: 149–162. doi:10.1016/S0960-9822(02)00448-7

33. Gordo I, Charlesworth B. The speed of Muller’s ratchet with background selection, and the degeneration of Y chromosomes. Genetical research. 2001;78: 149–161.

34. Charlesworth B, Charlesworth D. Rapid fixation of deleterious alleles can be caused by Muller’s ratchet. Genetical Research. 1997;70: 63–73. doi:10.1017/S0016672397002899

35. Engelstädter J. Muller’s ratchet and the degeneration of Y chromosomes: A simulation study. Genetics. 2008;180: 957–967. doi:10.1534/genetics.108.092379

36. Grossen C, Neuenschwander S, Perrin N. The evolution of XY recombination: Sexually antagonistic selection versus deleterious mutation load. Evolution. 2012;66: 3155–3166. doi:10.1111/j.1558-5646.2012.01661.x

37. Blaser O, Grossen C, Neuenschwander S, Perrin N. Sex-chromosome turnovers induced by deleterious mutation load. Evolution. 2013;67: 635–645. doi:10.1111/j.1558-5646.2012.01810.x

38. Blaser O, Neuenschwander S, Perrin N. Sex-chromosome turnovers: The hot-potato model. The American Naturalist. 2014;183: 140–146. doi:10.1086/674026

39. Lenormand T, Fyon F, Sun E, Roze D. Sex chromosome degeneration by regulatory evolution. Current Biology. 2020;30: 3001–3006.e5. doi:10.1016/j.cub.2020.05.052

40. Manna F, Martin G, Lenormand T. Fitness landscape: an alternative theory for the dominance of mutations. Genetics. 2011;189: 923–937.

41. Charlesworth B, Betancourt AJ, Kaiser VB, Gordo I. Genetic recombination and molecular evolution. Cold Spring Harbor Symposia on Quantitative Biology. 2009;74: 177–186. doi:10.1101/sqb.2009.74.015

42. Kaiser VB, Charlesworth B. Muller’s ratchet and the degeneration of the *Drosophila miranda* neo-*Y* chromosome. Genetics. 2010;185: 339–348. doi:10.1534/genetics.109.112789

43. Olito C, Abbott JK. The evolution of suppressed recombination between sex chromosomes and the lengths of evolutionary strata. Evolution. 2023; qpad023. doi:10.1093/evolut/qpad023

44. Porubsky D, Höps W, Ashraf H, Hsieh P, Rodriguez-Martin B, Yilmaz F, et al. Recurrent inversion polymorphisms in humans associate with genetic instability and genomic disorders. Cell. 2022;185: 1986–2005.e26. doi:10.1016/j.cell.2022.04.017

45. Cáceres M, Puig M, Ruiz A. Molecular characterization of two natural hotspots in the *Drosophila buzzatii* genome induced by transposon insertions. Genome Res. 2001;11: 1353–1364. doi:10.1101/gr.174001

46. Feuk L, MacDonald JR, Tang T, Carson AR, Li M, Rao G, et al. Discovery of human inversion polymorphisms by comparative analysis of human and chimpanzee DNA sequence assemblies. PLoS Genet. 2005;1: e56. doi:10.1371/journal.pgen.0010056

47. Dobzhansky TH. Genetics of the evolutionary process. New York: Columbia University Press; 1970.

48. Porubsky D, Höps W, Ashraf H, Hsieh P, Rodriguez-Martin B, Yilmaz F, et al. Recurrent inversion polymorphisms in humans associate with genetic instability and genomic disorders. Cell. 2022;185: 1986–2005.e26. doi:10.1016/j.cell.2022.04.017

49. Porubsky D, Sanders AD, Höps W, Hsieh P, Sulovari A, Li R, et al. Recurrent inversion toggling and great ape genome evolution. Nat Genet. 2020;52: 849–858. doi:10.1038/s41588-020-0646-x

50. Giner-Delgado C, Villatoro S, Lerga-Jaso J, Gayà-Vidal M, Oliva M, Castellano D, et al. Evolutionary and functional impact of common polymorphic inversions in the human genome. Nat Commun. 2019;10: 4222. doi:10.1038/s41467-019-12173-x

51. Lemaitre C, Braga MDV, Gautier C, Sagot M-F, Tannier E, Marais GAB. Footprints of inversions at present and past pseudoautosomal boundaries in human sex chromosomes. Genome Biology and Evolution. 2009;1: 56–66. doi:10.1093/gbe/evp006

52. Zhang W, Wai CM, Ming R, Yu Q, Jiang J. Integration of genetic and cytological maps and development of a pachytene chromosome-based karyotype in papaya. Tropical Plant Biol. 2010;3: 166–170. doi:10.1007/s12042-010-9053-2

53. Peichel CL, McCann SR, Ross JA, Naftaly AFS, Urton JR, Cech JN, et al. Assembly of the threespine stickleback Y chromosome reveals convergent signatures of sex chromosome evolution. Genome Biology. 2020;21: 1–31. doi:10.1186/s13059-020-02097-x

54. Navarro A, Betrán E, Barbadilla A, Ruiz A. Recombination and gene flux caused by gene conversion and crossing over in inversion heterokaryotypes. Genetics. 1997;146: 695–709. doi:10.1093/genetics/146.2.695

55. Koury SA. Predicting recombination suppression outside chromosomal inversions in Drosophila melanogaster using crossover interference theory. Heredity (Edinb). 2023. doi:10.1038/s41437-023-00593-x

56. Pegueroles C, Ordóñez V, Mestres F, Pascual M. Recombination and selection in the maintenance of the adaptive value of inversions. Journal of Evolutionary Biology. 2010;23: 2709–2717. doi:10.1111/j.1420-9101.2010.02136.x

57. Stevison LS, Hoehn KB, Noor MAF. Effects of inversions on within- and between-species recombination and divergence. Genome Biology and Evolution. 2011;3: 830–841. doi:10.1093/gbe/evr081

58. Levine RP. Crossing over and inversions in coadapted systems. The American Naturalist. 1956 [cited 6 Feb 2023]. doi:10.1086/281905

59. Ishii T, Nagai A, Hirono T, Nakamura K. Hibernation of mosquitoes in rock caves on Miyato island. Sci Rep Tôhoku Univ Ser IV. 1964;30: 159–165.

60. Korunes KL, Noor MAF. Pervasive gene conversion in chromosomal inversion heterozygotes. Mol Ecol. 2019;28: 1302–1315. doi:10.1111/mec.14921

61. Kupiec M, Petes TD. Allelic and ectopic recombination between Ty elements in yeast. Genetics. 1988;119: 549–559. doi:10.1093/genetics/119.3.549

62. Montgomery EA, Huang SM, Langley CH, Judd BH. Chromosome rearrangement by ectopic recombination in Drosophila melanogaster : genome structure and evolution. Genetics. 1991;129: 1085–1098. doi:10.1093/genetics/129.4.1085

63. Lambert S, Mizuno K, Blaisonneau J, Martineau S, Chanet R, Fréon K, et al. Homologous recombination restarts blocked replication forks at the expense of genome rearrangements by template exchange. Molecular Cell. 2010;39: 346–359. doi:10.1016/j.molcel.2010.07.015

64. Li W-H, Gu Z, Cavalcanti ARO, Nekrutenko A. Detection of gene duplications and block duplications in eukaryotic genomes. J Struct Funct Genomics. 2003;3: 27–34.

65. Watanabe Y, Takahashi A, Itoh M, Takano-Shimizu T. Molecular spectrum of spontaneous de novo mutations in male and female germline cells of *Drosophila melanogaster*. Genetics. 2009;181: 1035–1043. doi:10.1534/genetics.108.093385

66. Turner DJ, Miretti M, Rajan D, Fiegler H, Carter NP, Blayney ML, et al. The rates of de novo meiotic deletions and duplications causing several genomic disorders in the male germline. Nat Genet. 2008;40: 90–95. doi:10.1038/ng.2007.40

67. Lipinski KJ, Farslow JC, Fitzpatrick KA, Lynch M, Katju V, Bergthorsson U. High spontaneous rate of gene duplication in *Caenorhabditis elegans*. Curr Biol. 2011;21: 306–310. doi:10.1016/j.cub.2011.01.026

68. Charlesworth D, Bergero R, Graham C, Gardner J, Keegan K. How did the guppy Y chromosome evolve? PLOS Genetics. 2021;17: e1009704. doi:10.1371/journal.pgen.1009704

69. Sharma A, Heinze SD, Wu Y, Kohlbrenner T, Morilla I, Brunner C, et al. Male sex in houseflies is determined by Mdmd, a paralog of the generic splice factor gene CWC22. Science. 2017;356: 642–645. doi:10.1126/science.aam5498

70. Meisel RP, Olafson PU, Adhikari K, Guerrero FD, Konganti K, Benoit JB. Sex chromosome evolution in Muscid flies. G3 Genes|Genomes|Genetics. 2020;10: 1341–1352. doi:10.1534/g3.119.400923

71. Furman BLS, Metzger DCH, Darolti I, Wright AE, Sandkam BA, Almeida P, et al. Sex chromosome evolution: So many exceptions to the rules. Genome Biology and Evolution. 2020;12: 750–763. doi:10.1093/gbe/evaa081

72. Beukeboom LW, Perrin N. The Evolution of Sex Determination. Oxford University Press; 2014. doi:10.1093/acprof:oso/9780199657148.001.0001

73. Rodrigues N, Studer T, Dufresnes C, Perrin N. Sex-chromosome recombination in common frogs brings water to the Fountain-of-Youth. Molecular Biology and Evolution. 2018;35: 942–948. doi:10.1093/molbev/msy008

74. Perrin N. Sex reversal: a fountain of youth for sex chromosomes? Evolution. 2009;63: 3043–3049. doi:10.1111/j.1558-5646.2009.00837.x

75. Krimbas CB, Powell JR. Drosophila inversion polymorphism. CRC Press; 1992.

76. Hinton T. A correlation of phenotypic changes and chromosomal rearrangements at the two ends of an inversion. Genetics. 1950;35: 188–205. doi:10.1093/genetics/35.2.188

77. Novitski E. The regular reinversion of the roughest3 inversion. Genetics. 1961;46: 711–717. doi:10.1093/genetics/46.7.711

78. Emmens CW. Salivary gland cytology of Roughest3 inversion and reinversion, and Roughest2. Journ of Genetics. 1937;34: 191–202. doi:10.1007/BF02982262

79. Grüneberg H. The position effect proved by a spontaneous reinversion of the X-chromosome in *Drosophila melanogaster*. Journ of Genetics. 1937;34: 169–189. doi:10.1007/BF02982261

80. Muyle A, Marais G, Bačovský V, Hobza R, Lenormand T. Dosage compensation evolution in plants: theories, controversies and mechanisms. Philosophical Transactions of the Royal Society B. 2022;377: 20210222.

81. Gu L, Walters JR. Evolution of sex chromosome dosage compensation in animals: A beautiful theory, undermined by facts and bedeviled by details. Genome Biology and Evolution. 2017;9: 2461–2476. doi:10.1093/gbe/evx154

82. Tezenas E, Giraud T, Véber A, Billiard S. The fate of recessive deleterious or overdominant mutations near mating-type loci under partial selfing. Peer Community Journal. 2023;3: e14. doi:10.24072/pcjournal.238

